# Effects of hyperbaric environment on endurance and metabolism are exposure time-dependent in well-trained mice

**DOI:** 10.1101/2020.08.26.268995

**Authors:** Junichi Suzuki

## Abstract

Hyperbaric exposure (1.3 atmospheres absolute with 20.9% O_2_) for 1 h a day was shown to improve exercise capacity. The present study was designed to reveal whether the daily exposure time affects exercise performance and metabolism in skeletal and cardiac muscles. Male mice in the training group were housed in a cage with a wheel activity device for 7 weeks from 5 weeks old. Trained mice were then subjected to hybrid training (HT, endurance exercise for 30 min followed by sprint interval exercise for 30 min). Hyperbaric exposure was applied following daily HT for 15 min (15HT), 30 min (30HT), or 60 min (60HT) for 4 weeks (each group, n = 10). In the endurance capacity test, maximal work values were significantly increased by 30HT and 60HT. In the left ventricle (LV), activity levels of 3-hydroxyacyl-CoA-dehydrogenase, citrate synthase, and carnitine palmitoyl transferase (CPT) 2 were significantly increased by 60HT. CPT2 activity levels were markedly increased by 60HT in the plantaris muscle (PL). Pyruvate dehydrogenase complex (PDHc) activity values in PL were enhanced more by 30HT and 60HT than by HT. In both 30HT and 60HT groups, lactate dehydrogenase activity levels were significantly increased and decreased, respectively, in the gastrocnemius muscle and LV. These results indicate that hyperbaric exposure for 30 min or longer has beneficial effects on endurance, and 60-min exposure has the potential to further increase performance by facilitating fatty acid metabolism in skeletal and cardiac muscles in highly-trained mice.

## Introduction

Athletes with a broad range of performance levels use a commercial hyperbaric apparatus, which functions at <1.5 atmospheres absolute (ATA) with room air (20.9% O_2_). Acute hyperbaric exposure (1.3 ATA) for 1 h markedly up-regulated mRNA expression levels of proliferator-activated receptor gamma coactivator 1-alpha (PGC-1α) and peroxisome proliferator-activated receptor alpha (PPARα) in hind-leg muscles 3 h after exposure [1]. Moreover, endurance exercise training followed by hyperbaric exposure for daily 1 h markedly improved both fatty acid and glucose metabolism in hind-leg muscles in highly trained mice [2]. To apply daily hyperbaric exposure to human athletes, the duration of exposure should be shorter, and should not disturb the daily training regimen. Acute hyperbaric exposure (1.3 ATA with room air) for 30 min after maximal exercise markedly reduced the lactate concentration and heart rate in humans [3]. However, an additive effect of hyperbaric exposure on exercise performance has not yet been reported in terms of the daily exposure time.

High-intensity interval exercise (6 × 30-sec all-out cycling) followed by endurance exercise (60% VO_2_max for 60 min) markedly enhanced mRNA levels of PGC-1α compared with those observed after respective, single, uncombined regimens in trained human muscle [4]. Hybrid exercise training (HT) consisting of interval and endurance exercise may be beneficial to improve the endurance capacity of highly-trained individuals. HT (endurance exercise for 30 min followed by sprint interval exercise (5-s run-10-s rest) for 30 min) for 4 weeks enhanced the endurance capacity, but endurance training did not, in mice that were trained from an early age [5].

Although hyperbaric exposure for 1 h markedly facilitated exercise-induced muscle hypertrophy in mice [2], mechanisms underlying these changes have yet to be clearly determined. In previous studies, expression levels of heat shock protein (HSP) 70 were upregulated by endurance training with hyperbaric exposure in hind-leg muscles [1, 2]. HSP70 was shown to have an important role in the recovery of skeletal muscle after intensive exercise [6]. AKT, also known as protein kinase B (PKB), is a serine/threonine protein kinase with multiple regulatory functions, including the control of cell growth, survival, apoptosis, proliferation, angiogenesis, and the metabolism of carbohydrates, lipids and proteins [7]. AKT1 was shown to have a crucial role in regulating satellite cell proliferation during skeletal muscle hypertrophy [8].

Expression levels of mitochondrial fusion and fission proteins were shown to be affected by exercise with hyperbaric exposure [2]. In highly-trained mice, mitochondrial fission marker dynamin-related protein (DRP)-1, but not mitofusion (MFN)-1, expression levels were upregulated after endurance exercise training under normobaric conditions with hyperbaric exposure for 1 h daily [2]. However, the effects of HT with different durations of hyperbaric exposure on expression levels of these proteins remain unknown.

Most ATP production in the heart depends on fatty acid oxidation [9]. Fat metabolism in the heart was shown to decrease after chronic endurance training [10] and interval training [11]. However, no study has observed enzyme activity levels concerning fatty acid metabolism after chronic exercise training with hyperbaric exposure.

In the present study, experiments were designed to elucidate the effects of HT with different durations of hyperbaric exposure on exercise capacity as well as the expression of proteins involved in muscle growth and mitochondrial biogenesis, and metabolic enzyme activity levels in skeletal and cardiac muscles of well-trained mice.

## Materials and methods

### Ethical approval

All procedures were approved by the Animal Care and Use Committee of Hokkaido University of Education (No. 3, approved on 2019/4/16) and performed in accordance with the "Guiding Principles for the Care and Use of Animals in the Field of Physiological Sciences" of the Physiological Society of Japan and the 'European Convention for the Protection of Vertebrate Animals used for Experimental and other Scientific Purposes' (Council of Europe No. 123, Strasbourg, 1985).

### Animals, hyperbaric exposure, and exercise training

Fifty male ICR (MCH) mice (four weeks old) were purchased from Clea Japan Inc. (Tokyo, Japan) and housed under the conditions of a controlled temperature (24±1°C) and relative humidity of approximately 50%. Lighting (7:00-19:00) was controlled automatically. All mice were given commercial laboratory chow (solid CE-2, Clea Japan) and tap water *ad libitum*. After mice had been fed for one week and allowed to adapt to the new environment, they were randomly assigned to a sedentary control group (Sed, n=10) or training group (n=40). Mice in the training group were individually housed in a cage with a wheel activity device (13 cm in diameter) for 7 weeks, as described previously [2]. Wheel activity (distance and running time) was monitored and recorded using digital bike computers (CC-VL820, Cateye Co., Ltd., Osaka, Japan). Mice in the Sed group were housed individually throughout the experiment. To familiarize mice with the treadmill device, all mice including the Sed group were subjected to treadmill walking once a week using a controlled treadmill (Modular motor assay, Columbus Instruments Inc., Columbus, OH, USA) for 5 min per day at 10-15 m min^-1^ with a 5-deg incline.

Running distance during voluntary wheel training is shown in Fig 1. Following voluntary wheel training, mice were given a 48-h non-exercise period prior to the maximal endurance capacity test. The test was performed with a graded ramp running protocol using the controlled treadmill, as described previously [5]. Total work (joules) was calculated as a product of body weight (kg), speed (m sec^-1^), time (sec), slope (%), and 9.8 (m sec^-2^). Exhaustion was defined when the mouse stayed for more than 5 s on the metal grid (no electrical shock) at the rear of the treadmill, despite external gentle touch being applied to their tail with a conventional elastic bamboo stick (0.8 mm in diameter).

**Fig 1.**
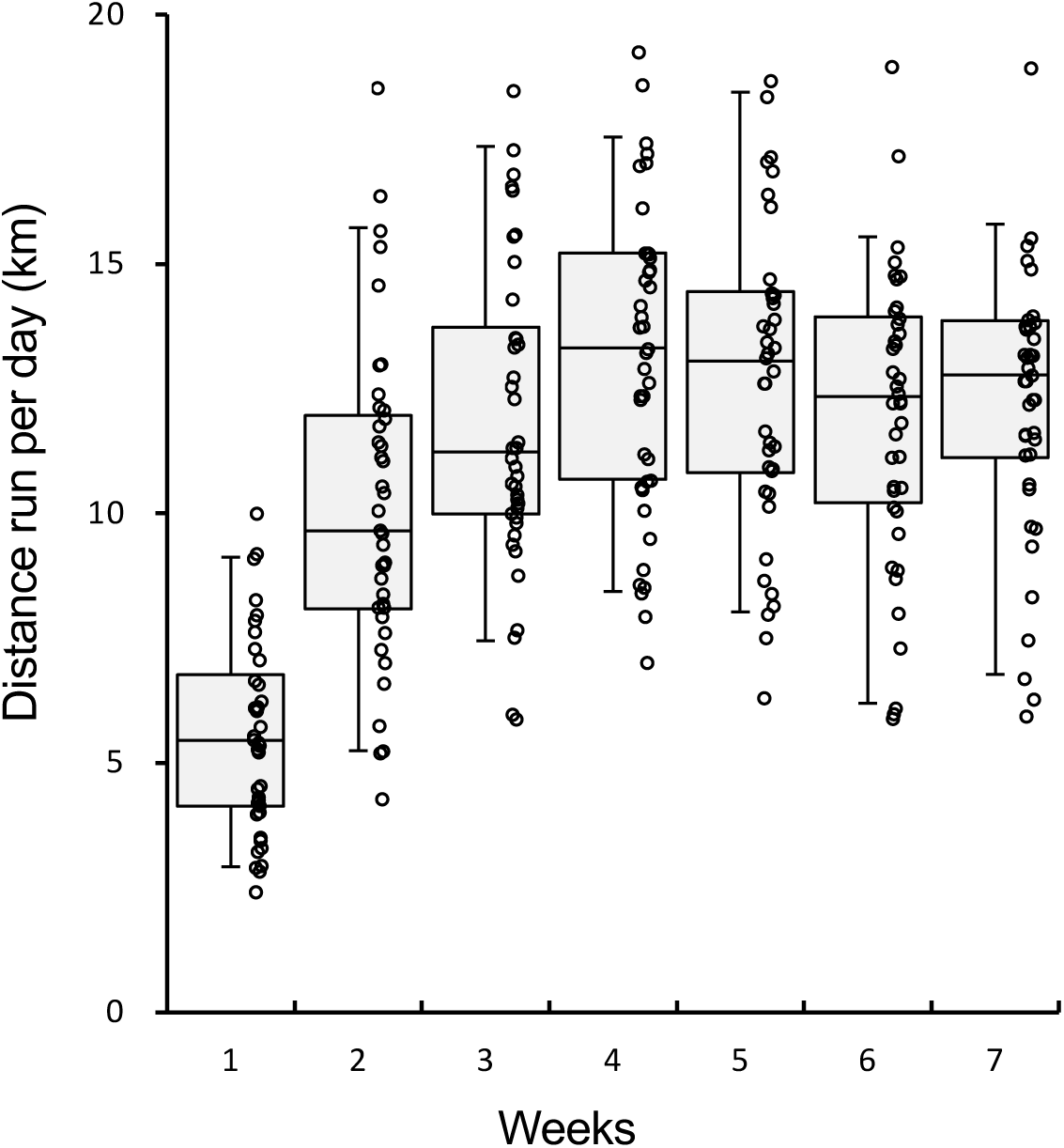
Running distance per day during voluntary wheel training. Values are expressed as box and whisker plots with 5th, 25th, 50th, 75th and 95th percentile. Dots are individual data points.

Following the performance test, mice were given a 48-h non-exercise period prior to treadmill training. Mice in the training group were divided into a hybrid-training group (HT, n=10), hybrid-training group with hyperbaric exposure for 15 min daily (15HT, n=10), 30 min daily (30HT, n=10), or 60 min daily (60HT, n=10), in order to match the mean and SD values of total work (Fig 2A). Mice in the training groups were subjected to HT in a normobaric environment 6 days per week for 4 weeks (within 2 hours from approximately 5 AM) using a rodent treadmill (KN-73, Natsume Co., Tokyo, Japan). Instead of using an electrical shock, the tali or feet of mice were touched with a conventional test tube brush made of soft porcine bristles in order to motivate them to run when they stayed on a metal grid for more than 2 s. HT consists of endurance exercise for 30 min followed by an interval exercise regimen (5-sec run-10-sec rest) for 30 min interposed by a 5-min rest. For endurance exercise, mice ran at 20 m min^-1^ with a 15-deg incline on the first and second days of training. The running speed was increased to 25 and 27.5 m min^-1^ on the first and fourth days of the second week, respectively, and to 30 m min^-1^ on the second day of the third week. For the interval exercise regimen, mice ran at 30 m min^-1^ with a 15-deg incline on the first day of training. The running speed was increased to 35, 37.5, 40, and 42.5 m min^-1^ on the 4th, 7th, 10th, and 13th days of training, respectively. The maximal endurance capacity test was performed 48 hours after the last run, as described above.

**Fig 2.**
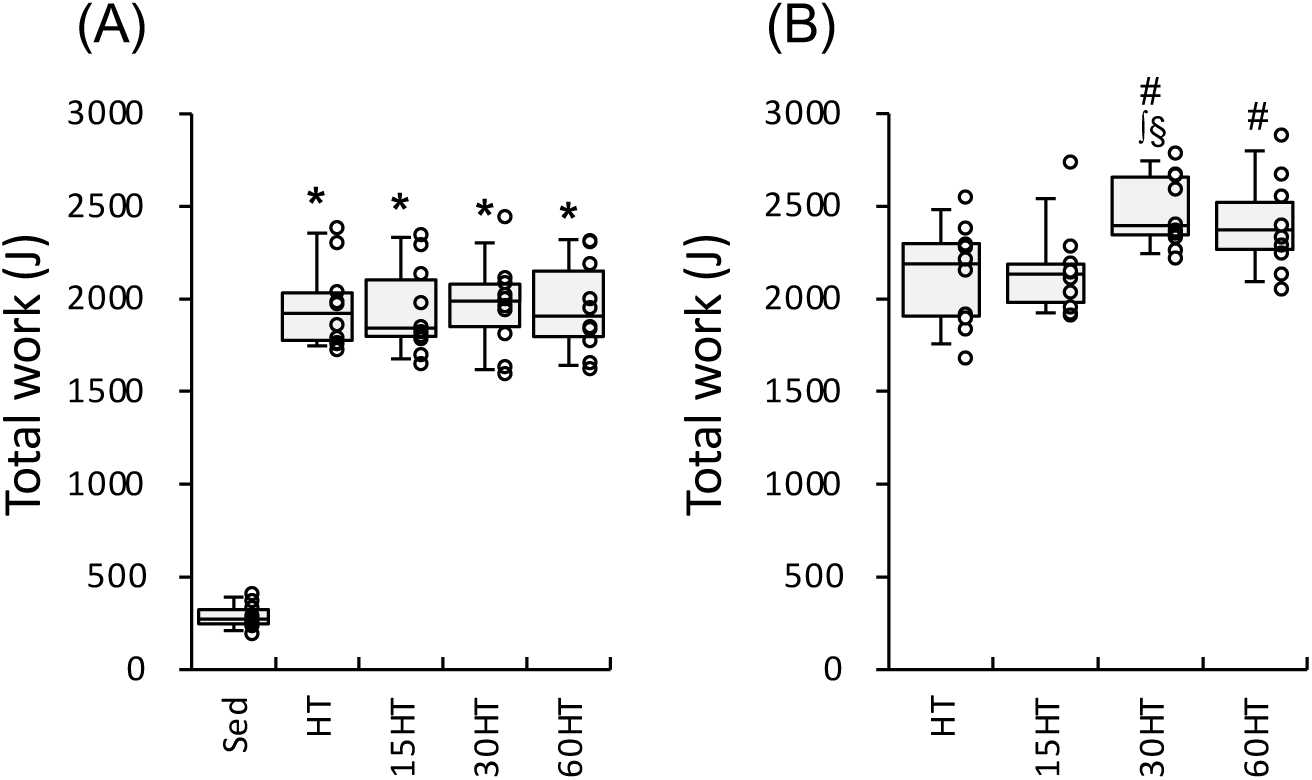
Endurance exercise performance test. Total work capacity of the endurance capacity test after 7 weeks of voluntary wheel running (A) and after 4 weeks of treadmill exercise training (B). #, significantly different from pre-treadmill training values of each group shown in the panel A. *, ∫, and §, significantly different from Sed, HT, and 15HT groups, respectively. Values are expressed as box and whisker plots with 5th, 25th, 50th, 75th and 95th percentile. Dots are individual data points.

The hyperbaric exposure group was subjected to 1.3 ATA with room air (20.9% O_2_) for 15, 30, or 60 min, once per day (started at approximately 30 min after daily exercise), 6 days per week, for 4 weeks. The hyperbaric exposure was performed as described previously [2]. The pressure of the chamber was gradually increased or decreased at 0.136 ATA min^-1^ within 2.2 min.

Forty-eight hours after the last run, mice were anesthetized with α-chloralose (0.06 g kg^-1^ i. p. Wako Pure Chemical Industries Ltd., Osaka, Japan) and urethane (0.7 g kg^-1^ i. p. Wako). A toe pinch response was used to validate adequate anesthesia. The plantaris (PL), and gastrocnemius muscles were excised and the deep red region (Gr) of the gastrocnemius was isolated from the superficial white region (Gw). The diaphragm (DIA) was excised. All samples were frozen in liquid nitrogen for biochemical analyses. The remaining muscles, i.e., those on the right side, were excised and placed in embedding medium, O.C.T. compound (Miles Inc., Elkhart, IN, USA), and then rapidly frozen in isopentane cooled to its melting point (-160°C) with liquid nitrogen. Mice were killed by excision of the heart. The whole heart and left ventricle (LV) were weighed and frozen in liquid nitrogen. All tissue samples were stored at −80°C until for analyses.

### Histological analyses

Histochemical examinations of capillary profiles and muscle fiber phenotypes were conducted as previously reported by the author with slight modifications [5]. Briefly, ten-micrometer-thick serial cross-sections were obtained using a cryotome (CM-1500; Leica Japan Inc., Tokyo, Japan) at −20°C from the mid-belly portion of calf muscles. These sections were air-dried, fixed with 100% ethanol at 4°C for 15 min, and then washed in 0.1 M phosphate-buffered saline (PBS) with 0.1% Triton X-100. Sections were then blocked with 10% goat normal serum at room temperature for 30 min, washed in PBS for 5 min, and incubated at 4°C overnight with a mixture of fluorescein-labelled Griffonia simplicifolia lectin (GSL I) (FL 1101 (1:100), Vector Laboratories Inc., California, USA), an anti-type I myosin heavy chain (MHC) antibody (BA-F8, mouse IgG2b, 1:80), and anti-type IIA MHC antibody (SC-71, mouse IgG1, 1:80) diluted with PBS. Sections were then reacted with a secondary antibody mixture containing Alexa Fluor 350-labeled anti-mouse IgG2b (1:100) and Alexa Fluor 647-labeled anti-mouse IgG1 (1:100) diluted with PBS at room temperature for 1 h. Sections were coverslipped with Fluoromount/Plus (K048, Diagnostic BioSystems Co., California, USA). Primary and secondary antibodies were purchased from the Developmental Studies Hybridoma Bank (University of Iowa) and Thermo Fisher Scientific Inc. (Tokyo, Japan), respectively. Fluorescent images of the incubated sections were observed using a microscope (Axio Observer, Carl Zeiss Japan, Tokyo, Japan). Muscle fiber phenotypes were classified as type I (blue), type I+IIA (faint blue and faint red), type IIA (red), type IIAX (faint red), and type IIB+IIX (unstained). Non-overlapping microscopic fields were selected at random from each muscle sample. The observer was blinded to the source (groups) of each slide during the measurements. Fluorescent images were obtained from PL, the lateral (GrL) and medial (GrM) portions of Gr, and Gw. Fiber cross-sectional area (FCSA) values were obtained using public domain Image J software (NIH, Bethesda, MD, USA). The negative control without primary antibodies was confirmed to show no fluorescent signal.

### Biochemical analyses of enzyme activities

Frozen tissue powder was obtained using a frozen sample crusher (SK mill, Tokken Inc., Chiba, Japan) and homogenized with ice-cold medium (10 mM HEPES buffer, pH 7.4; 0.1% Triton X-100; 11.5% (w/v) sucrose; and 5% (v/v) protease inhibitor cocktail (P2714, Sigma-Aldrich)). After centrifugation at 1,500 × *g* at 4°C for 10 min, the supernatant was used in enzyme activity analyses. The activities of 3-hydroxyacyl-CoA-dehydrogenase (HAD) and lactate dehydrogenase (LDH) were assayed according to the method of Bass *et al.* [12]. The activities of citrate synthase (CS) and phosphofructokinase (PFK) were assayed according to the method of Srere [13] and Passonneau and Lowry [14], respectively. Pyruvate dehydrogenase complex (PDHc) activity was measured by the phenazine methosulfate-3-(4,5-dimethylthiazol-2-yl)-2,5-diphenyltetazolium bromide (PMS-MTT) assay [15]. For the carnitine palmitoyl transferase (CPT) 2 assay, the tissue homogenate was added to the reaction medium (200 mM HEPES, 10 mM EGTA, 0.2 M sucrose, 400 mM KCl, 2 mM DTNB (Sigma O4126), 0.13% (w/v) bovine serum albumin (Wako 017-15146), 20 *µ*M palmityl CoA (Sigma P 9716), 10 *µ*M malonyl CoA (Sigma M4263)) and incubated for 1 h in a dark chamber at 25 ºC. After confirming there was no non-specific reaction, the reaction was initiated by adding 15 mM L-carnitine (Sigma C0283). All measurements were conducted at 25°C with a spectrophotometer (U-2001, Hitachi Co., Tokyo, Japan) and enzyme activities were obtained as *µ*Mol hour ^-1^ mg protein^-1^. Total protein concentrations were measured using PRO-MEASURE protein measurement solution (iNtRON Biotechnology Inc., Gyeonggi-do, Korea).

### Western blot analyses

The tissue homogenates described above were used for Western blot analyses. A sample (50 *µ*g) was fractionated by SDS/PAGE on 12% (w/v) polyacrylamide gels (TGX StainFree FastCast gel, Bio-Rad Inc., CA, USA), exposed to UV for 1 min and total protein patterns were visualized using ChemiDoc MP Imager (Bio-Rad). The stain-free gel contains a trihalo compound which reacts with proteins during separation, rendering them detectable using UV exposure [16, 17]. Then, gels were electrophoretically transferred to a polyvinylidene fluoride membrane. The bands in each lane on the membrane were detected with ChemiDoc MP and the images were used for normalization process as described below. The blots were blocked with 3% (w/v) bovine serum albumin (protease and IgG free, 010-25783 Fujifilm Wako Pure Chemical, Osaka Japan), 1% (w/v) polyvinylpyrrolidone (PVP40, Sigma-Aldrich), and 0.3% (v/v) Tween-20 in PBS for 1 h, and then exposed to a specific primary antibody (1:500, Santa Cruz Biotechnology Inc., California, USA) against AKT1 (sc-5298), HSP70 (1:500, sc-66048), TFAM (sc-166965), MFN2 (sc-100560), FABP (sc-514208), or DRP1 (1:500, sc-271583) diluted in PBS with 0.05% Tween-20 for 1 h. After the blots had been incubated with a HRP-labeled mouse IgGκ light chain binding protein (1:5000, sc-516102, Santa Cruz), they were reacted with Clarity Western ECL substrate (Bio-Rad), Clarity Max Western ECL substrate (Bio Rad), or their mixture and the required proteins were detected with ChemiDoc MP. The densities of the specific bands were quantified using Image Lab software (Bio Rad) and normalized to the densities of all protein bands in each lane on the membrane [16, 17]. Then, the normalized densities of the bands were normalized again to the same sample that was run on every gel and transferred to every membrane.

### Statistical analyses

Differences between the two groups were examined using Bayesian estimation with a gamma prior distribution proposed by Kruschke [18]. The posterior distribution was obtained using the Markov chain Monte Carlo (MCMC) methods. Public domain R, RStudio, and JAGS programs were used for computing Bayesian inference. The MCMC chains were considered to show a stationary distribution when the Gelman-Rubin values (shrink factor) were less than 1.10 for all parameters. The significance of differences was evaluated by the 95% highest density interval (HDI) and a region of practical equivalence (ROPE) [19]. When the HDI value on the effect size fell outside of the ROPE set at −0.1 to 0.1, the difference was regarded as significant [19, 20]. Bayesian robust linear regression [18] was used to establish any correlation between the parameters observed in the present study. The regression was considered significant when HDI on the slope fell outside of the ROPE. Data are expressed as box and whisker plots with 5th, 25th, 50th, 75th and 95th percentile in the figures and means ± standard deviation (SD) in Table 1.

**Table 1.** Body and organ masses

|  | Sed (n=10) | HT (n=10) | 15HT (n=10) | 30HT (n=10) | 60HT (n=10) |
| --- | --- | --- | --- | --- | --- |
| Body mass (g) |  |  |  |  |  |
| Pre-HT | 38.3±1.7 | 36.1±3.4 | 36.3±0.95 | 35.0±1.7 | 36.2±1.1 |
| Post-HT | 41.0±1.7 | 37.5±2.0 * | 38.1±0.9 * | 38.4±1.7 * | 38.4±1.2 * |
| Post-Pre difference | 2.71±1.16 | 1.45±1.77 | 1.81±1.12 | 3.35±1.15 | 2.19±1.17 |
| Organ mass (mg) |  |  |  |  |  |
| Gastrocnemius | 178.6±12.1 | 174.3±8.9 | 171.4±7.1 | 171.2±8.1 | 175.2±10.2 |
| Plantaris | 21.7±1.1 | 23.1±1.9 | 22.7±2.2 | 23.7±1.7 | 23.3±3.1 |
| Whole heart | 158.1±10.2 | 178.1±12.3 * | 182.6±9.6 * | 187.5±10.6 * | 183.3±7.6 * |
| Left ventricle | 114.3±8.9 | 128.4±9.9 * | 133.3±7.6 * | 136.2±8.7 * | 137.1±14.3 * |
| Organ mass-to-body mass ratio (mg g<sup>-1</sup>) |  |  |  |  |  |
| Gastrocnemius | 4.36±0.29 | 4.65±0.26 | 4.50±0.21 | 4.46±0.21 | 4.56±0.17 |
| Plantaris | 0.53±0.04 | 0.62±0.06 | 0.60±0.06 | 0.62±0.04 | 0.61±0.08 |
| Whole heart | 3.86±0.17 | 4.75±0.22 * | 4.79±0.18 * | 4.88±0.21 * | 4.78±0.12 * |
| Left ventricle | 2.79±0.15 | 3.42±0.16 * | 3.49±0.14 * | 3.55±0.18 * | 3.56±0.37 * |

Values are presented as means ± SD. \*, significantly different from the Sed group using Bayesian estimation. Pre-HT and Post-HT, at the beginning and end, respectively, of the 4 weeks of HT.

## Results

### Body and organ masses

Body weight values after HT were significantly lower in the four exercise-trained groups than in Sed group (Table 1, each group, n = 10). However, body weight values before HT and the amount of increase during HT were not significantly different among groups. The absolute and relative weight values of the whole heart and LV were significantly higher in the four exercise-trained groups than in the Sed group. The relative weight values of GAS and PL were higher in the four exercise-trained groups than in the Sed group, but the differences were not significant. Thus, hyperbaric exposure did not affect the body weight or HT-induced cardiac and skeletal muscle growth.

### Maximal exercise capacity

After voluntary wheel running for 7 weeks, total work values were significantly greater in the training group by 6.9-fold (Fig 2A) than in Sed group (each group, n = 10). In 30HT and 60HT groups, total work values were significantly increased after 4 weeks of treadmill training. Moreover, total work values were significantly greater in the 30HT group than in HT and 15HT groups. Thus, daily hyperbaric exposure longer than 30 min had additive effects on HT-induced improvements in endurance capacity.

### Metabolic enzyme activities

CPT2 activity values in Gr were significantly higher in the three exposure groups than in the Sed group and, moreover, the values were significantly higher in 15HT and 60HT groups than in the HT group (Fig 3C, each group, n = 10). In PL and LV, CPT2 activity levels showed significantly higher values in the 60HT group than in the Sed group. CPT2 activity values in Gw were significantly higher in 15HT and 30HT groups than in the Sed group. In DIA, CPT2 values were positively correlated with maximal work values (Table 2).

**Table 2.** Correlations

| Explanatory variable | Response variable | Gr | Gw | PL | DIA | LV |
| --- | --- | --- | --- | --- | --- | --- |
| LDH | maximal work ‡ | * | ns | ns | ns | ns |
| PDHc | maximal work ‡ | ns | ns | * | ns | ns |
| CPT2 | maximal work ‡ | ns | ns | ns | * | ns |
| CS | maximal work ‡ | ns | ns | ns | * | ns |
| AKT1 | Total FCSA | ns | ns | * | ns | ns |
| AKT1 | C:F ratio | ns | ns | * | ns | ns |
| HSP70 | Total FCSA | ns | * | ns | ns | ns |
| DRP1 | CS | * | * | * | * | * |
| DRP1 | HAD | * | * | * | * | * |
| DRP1 | CPT2 | ns | * | ns | ns | ns |
| TFAM | CS | * | * | ns | ns | * |
| TFAM | HAD | * | * | * | ns | * |
| MFN2 | CS | * | ns | ns | ns | ns |
| MFN2 | HAD | ns | ns | * | ns | ns |

Correlations were established using Bayesian robust linear regression. \*, significant correlation; ns, non-significant correlation. ‡, Correlations were calculated without Sed group.

**Fig 3.**
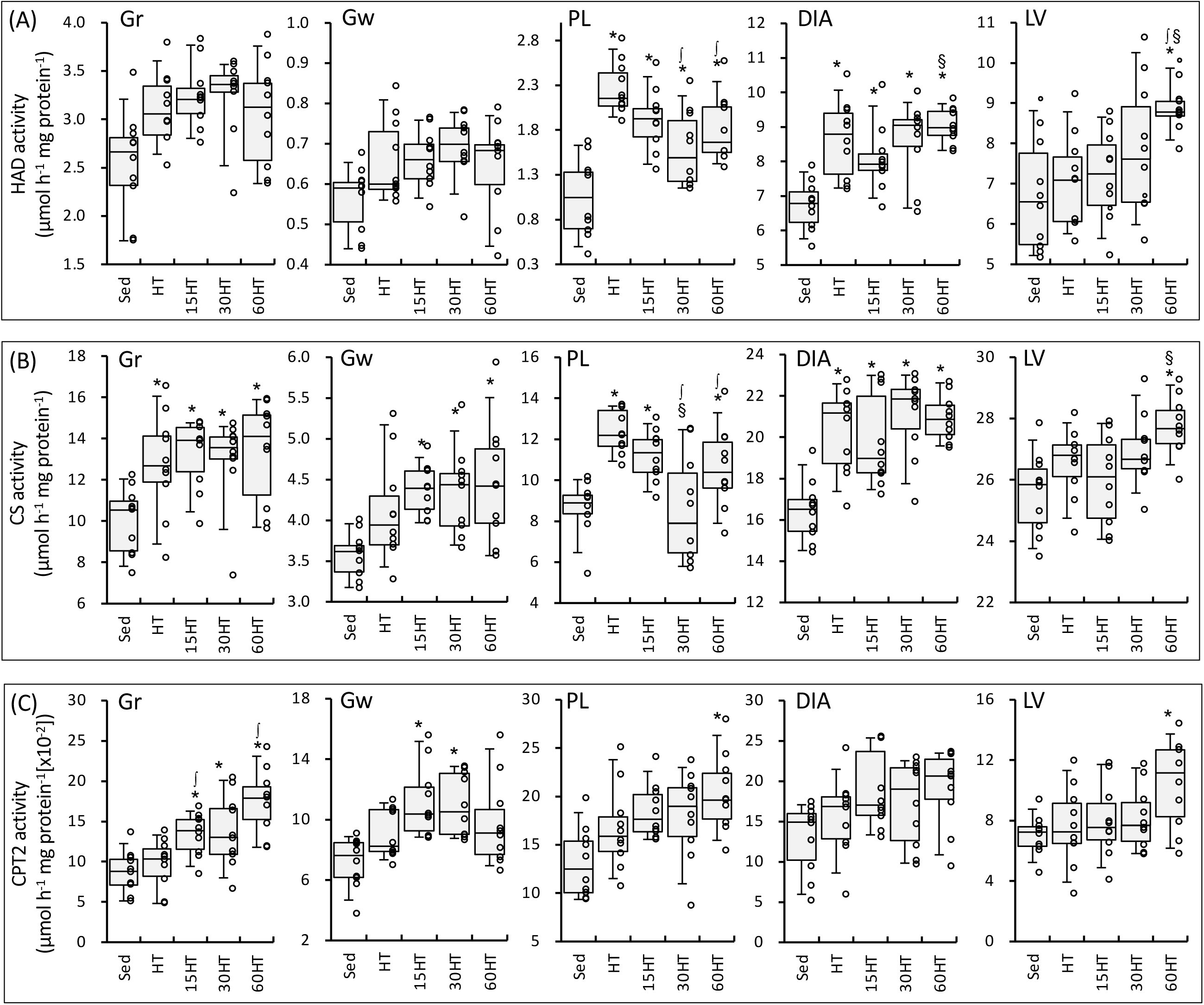
Enzyme activity values for HAD (A), CS (B), and CPT2 (C). Values are expressed as box and whisker plots with 5th, 25th, 50th, 75th and 95th percentile. Dots are individual data points. *, ∫, and §, significantly different from Sed, HT, and 15HT groups, respectively.

HAD activity values in LV were significantly higher in 60HT than in Sed, HT, and 15HT groups (Fig 3A, each group, n = 10). In PL, HAD levels were significantly lower in 30HT and 60HT groups than in the HT group, while activity values showed significantly greater values in the four training groups than in the Sed group.

CS activity values in LV were significantly higher in the 60HT than in Sed and 15HT groups (Fig 3B, each group, n = 10). In DIA, a positive correlation was observed between CS levels and maximal work values (Table 2).

PDHc activity values in PL were significantly higher in 30HT and 60HT groups than in Sed, HT, and 15HT groups (Fig 4A, each group, n = 10). Moreover, in PL, a positive correlation was found between PDHc levels and maximal work values (Table 2). In Gw, PDHc activity levels showed significantly greater values in 15HT, 30HT, and 60HT groups than in Sed and HT groups, and the levels were significantly greater in the 60HT group than in the 15HT group. In DIA, PDHc activity values were significantly higher in 15HT and 30HT groups than in the HT group. PDHc activity levels in LV showed significantly lower values in the 15HT group than in Sed and HT groups.

**Fig 4.**
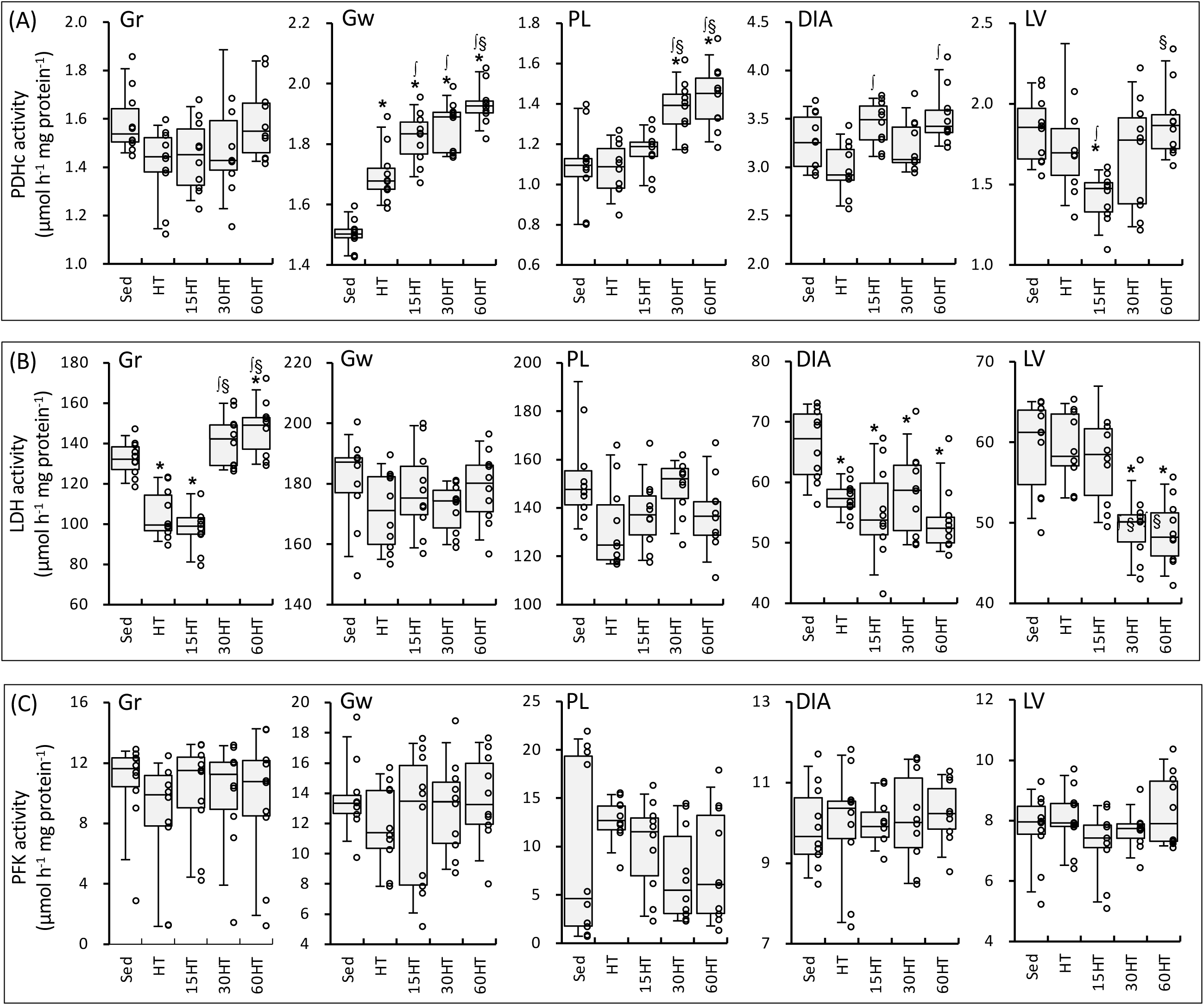
Enzyme activity values for PDHc (A), LDH (B), and PFK (C). Values are expressed as box and whisker plots with 5th, 25th, 50th, 75th and 95th percentile. Dots are individual data points. *, ∫, and §, significantly different from Sed, HT, and 15HT groups, respectively.

In 30HT and 60HT groups, LDH activity values were significantly higher and lower, respectively, in Gr and LV, than in Sed, HT, and 15HT groups (Fig 4B, each group, n = 10). LDH activity values were positively correlated with maximal work values in Gr (Table 2).

Thus, 60HT enhanced the activities of enzymes involved in mitochondrial fatty acid as well as glucose metabolism in hindleg and cardiac muscles, whereas 30HT predominantly enhanced glucose metabolism in the hindleg muscles.

### Protein expression

TFAM expression in Gr was significantly stronger in the four training groups than in the Sed group (Fig 5C, each group, n = 10). TFAM levels in PL were significantly higher in HT and 60HT groups than in the Sed group. MFN2 protein expression in PL was markedly stronger in the four training groups than in the Sed group (Fig 5B).

**Fig 5.**
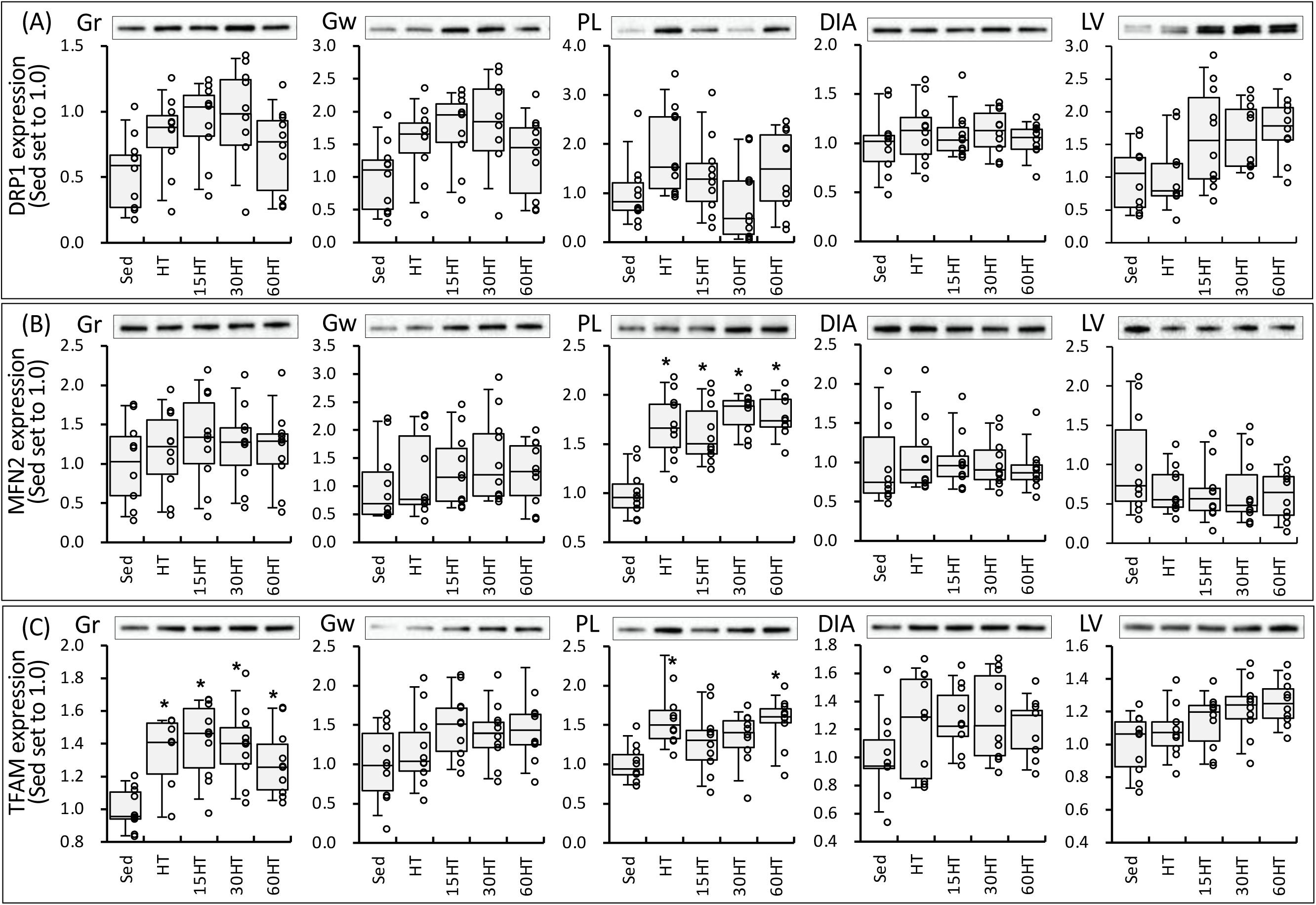
Expression of DRP1 (A), MFN2 (B), and TFAM (C) proteins. Values are expressed as box and whisker plots with 5th, 25th, 50th, 75th and 95th percentile. Dots are individual data points. *, significantly different from the Sed group.

AKT1 protein expression in PL was significantly stronger in the four training groups than in the Sed group (Fig 6B, each group, n = 10). AKT1 expression levels in PL were positively correlated with values of total FCSA and FCSA of all fiber types (Table 2). A significant positive correlation was observed between AKT1 expression levels and values in PL (Table 2).

**Figure 6.**
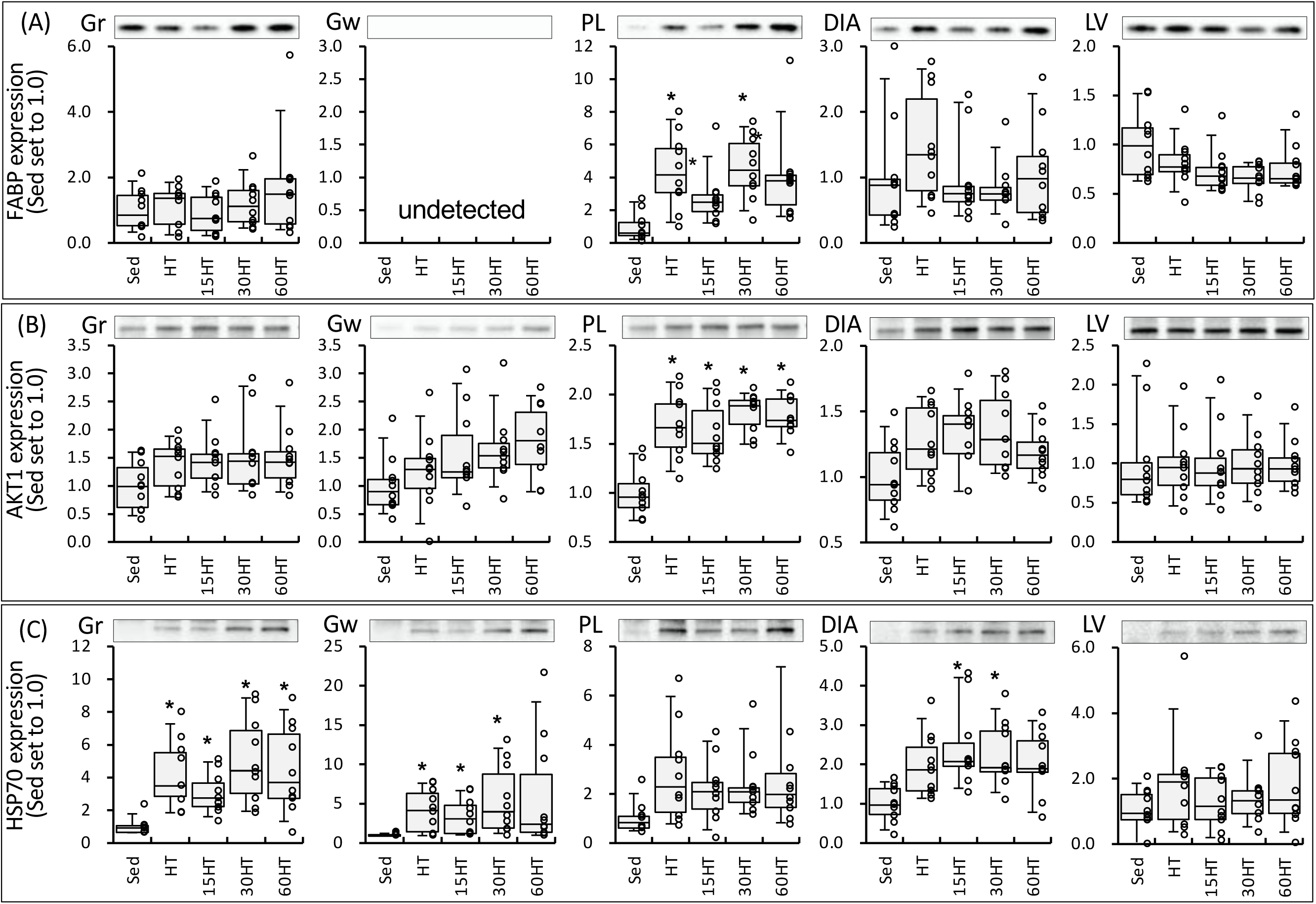
Expression of FABP (A), AKT1 (B), and HSP70 (C) proteins. Values are expressed as box and whisker plots with 5th, 25th, 50th, 75th and 95th percentile. Dots are individual data points. *, significantly different from the Sed group.

HSP70 expression levels in Gr were significantly stronger in the four training groups than in the Sed group (Fig 6C, each group, n = 10). In Gw, HSP70 protein expression was significantly stronger in HT, 15HT, and 30HT groups than in the Sed group. HSP70 levels in Gw tended to be greater (1.6-fold) in the 60HT group than in the Sed group. HSP70 expression levels in DIA were significantly stronger in 15HT and 30HT groups than in the Sed group. HSP70 levels were positively correlated with total FCSA values in Gw and with FCSA of all fiber types in PL (Table 2).

### Fiber type composition

HT with and without hyperbaric exposure significantly increased the proportion of type IIA and IIAX fibers and decreased the proportion of type IIB+IIX fibers in PL (Fig 7C, each group, n = 10). In GrL, the proportion of type IIAX fibers was larger and that of type IIB+IIX fibers was smaller in the four trained groups than in Sed group (Fig 7A). Thus, hyperbaric exposure did not affect the fiber proportion in hybrid-trained hindleg muscles.

**Fig 7.**
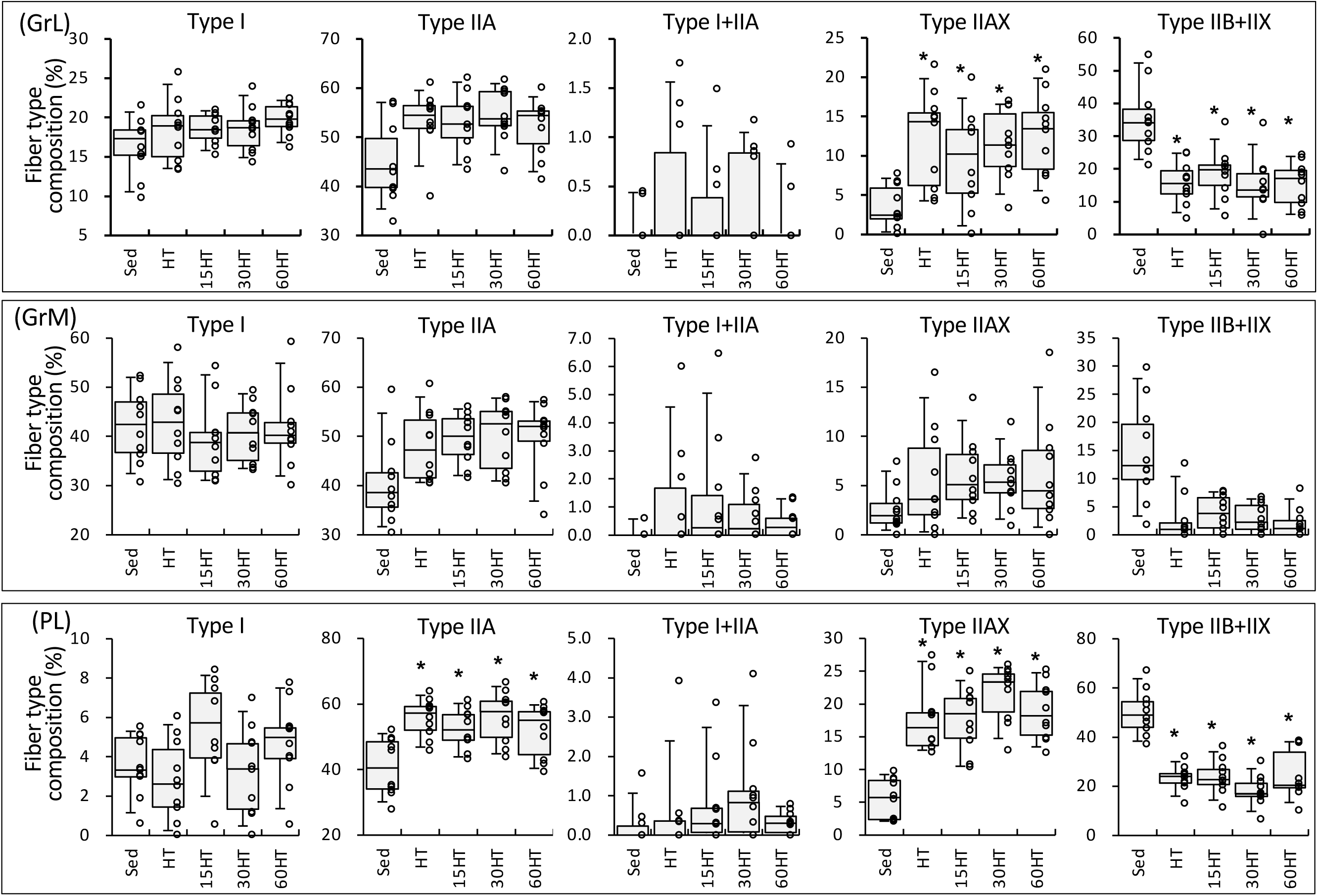
Fiber-type composition values in GrL (A), Grm (B), and PL (C) proteins. Values are expressed as box and whisker plots with 5th, 25th, 50th, 75th and 95th percentile. Dots are individual data points. *, significantly different from the Sed group.

### Fiber cross-sectional area

Total FCSA values in Gw were higher in the 60HT group than in Sed (by 16%) and 30HT (by 11%) groups, but the difference was not significant (Fig 8B). The distribution of total FCSA in the 60HT group was shifted to the larger fiber size in Gw (Fig 8A). Thus, hyperbaric exposure for 1 h partially promoted muscle fiber hypertrophy in highly glycolytic fibers.

**Fig 8.**
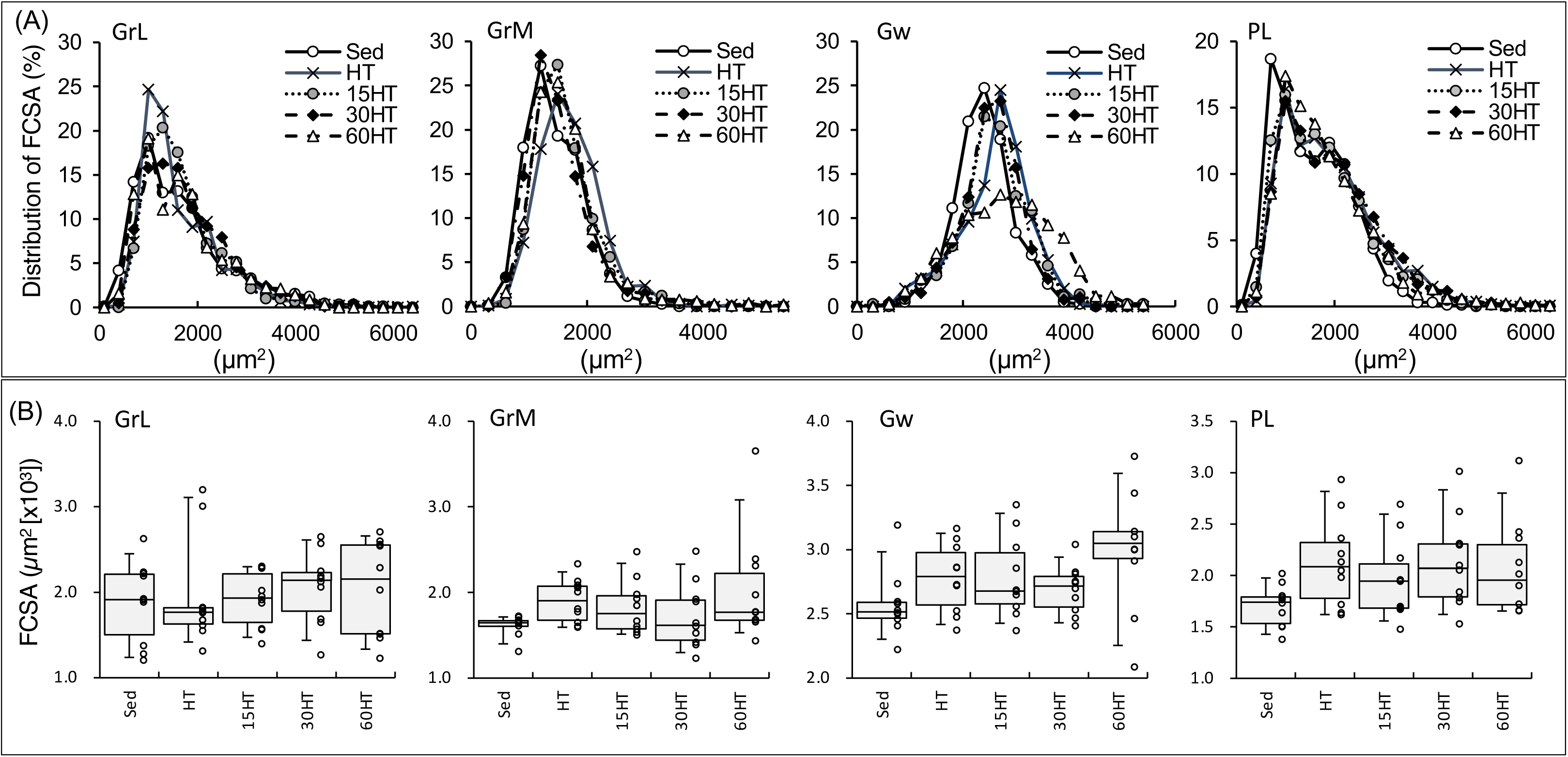
Distribution of FCSA (A) and total FCSA (B) values. Values are expressed as box and whisker plots with 5th, 25th, 50th, 75th and 95th percentile. Dots are individual data points in the panel (B).

### Capillarization

Capillary-to-fiber (C:F) ratio values in PL were significantly higher in the four trained groups than in the Sed group (Fig 9, each group, n = 10). C:F ratio values in GrL and GrM were higher in the four training groups, but the differences were not significant. C:F values in Gw were markedly higher in the 60HT group than in the Sed group by 21%. Thus, daily hyperbaric exposure for 1 h markedly facilitated exercise-induced capillary growth in the muscle regions mainly composed of glycolytic fibers.

**Fig 9.**
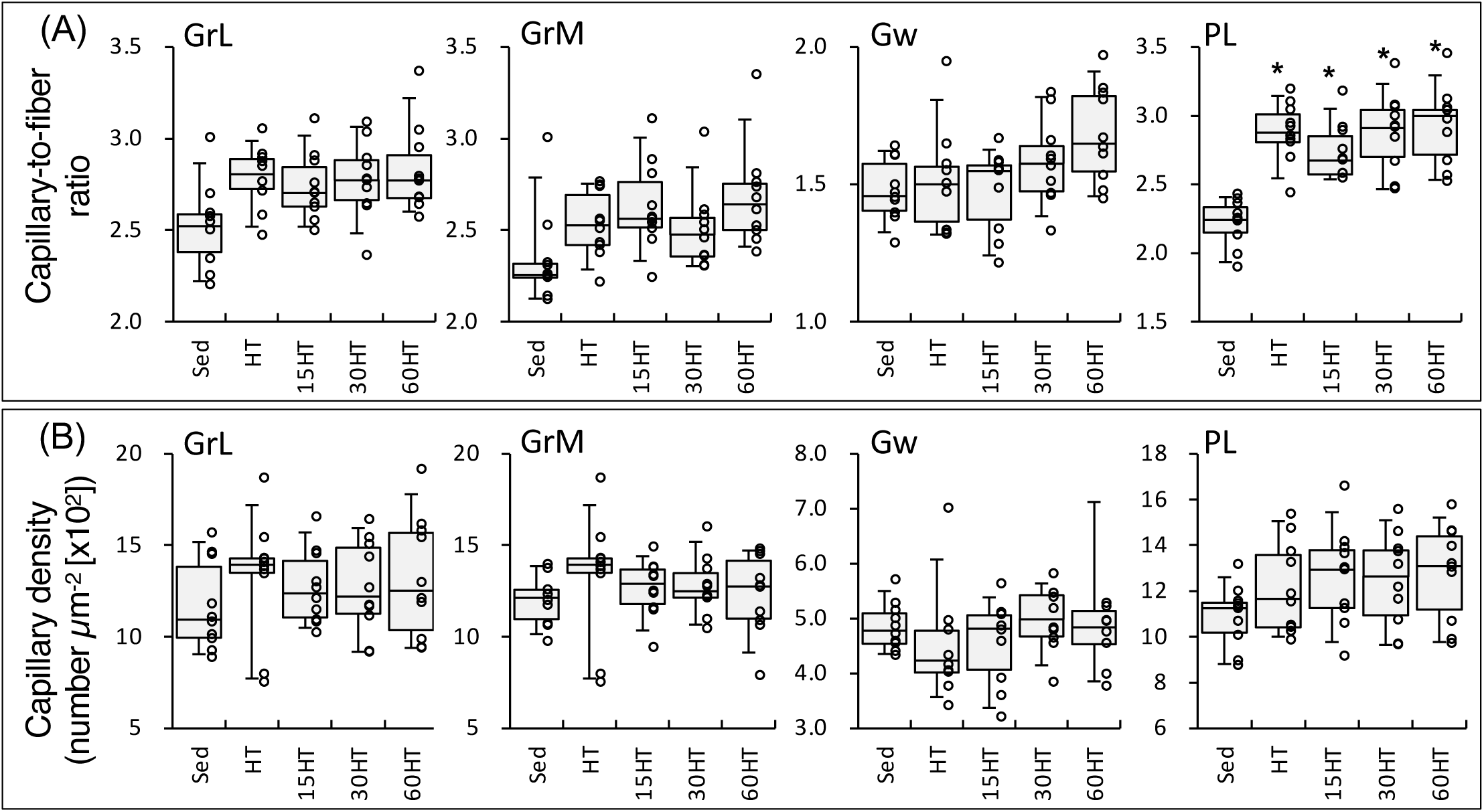
Capillary-to-fiber ratio (A) and capillary density (B) values. Values are expressed as box and whisker plots with 5th, 25th, 50th, 75th and 95th percentile. Dots are individual data points. *, significantly different from the Sed group.

## Discussion

In highly-trained mice, the present results showed that endurance exercise performance was promoted by HT with daily hyperbaric exposure for 30 and 60 min, but not for 15 min. In both 30HT and 60HT groups, additive effects of muscle metabolism were noted in PDHc activity values in PL and Gw, and in LDH activity values in Gr (Figs 3 and 4). Thus, daily hyperbaric exposure for 30-60 min had additive effects on the training-induced increase in glucose metabolism in hindleg muscles. Endurance training with 60 min of hyperbaric exposure daily was shown to enhance PFK activity values, but did not change LDH activity values in hindleg muscle of highly-trained mice [2], suggesting that HT with hyperbaric exposure for at least 30 min daily may be beneficial to improve glucose metabolism in hindleg muscles.

In addition to these beneficial changes, HT with daily hyperbaric exposure for 60 min markedly enhanced cardiac fatty acid metabolism, identified by marked increases in HAD, CS, and CPT2 activity levels in LV (Fig 3). The present study is the first to report the effects of exercise training with hyperbaric exposure on cardiac metabolism. Around 95% of ATP used by cardiomyocytes is generated by oxidative phosphorylation in mitochondria [9]. Up to 70% of the oxidative metabolism principally relies on fatty acid oxidation [9]. However, fatty acid metabolism in the heart was not shown to be enhanced by chronic exercise training. After 5-6 weeks of endurance training, palmitoylcarnitine oxidation was shown to be significantly reduced in the heart, whereas it was unchanged in the gastrocnemius muscle of rats [10]. Endurance training for 8 weeks did not increase palmitate oxidation while interval training (4-min run-2-min rest) decreased it in the mouse heart [11]. Consistent with these findings, in the present study, HT itself did not affect enzyme levels concerning fatty acid metabolism in LV.

Glycolytic energy production by active skeletal muscles increases in proportion to exercise intensity, thereby increasing the plasma lactate concentration [21]. Myocardial lactate oxidation is prominent and proportional to the exogenous concentration during elevated workloads in rats [22]. In isolated working rat hearts, lactate was found to contribute a relatively large proportion (up to 37%) of the total ATP supply under a low-fat condition [23]. In contrast, under a high-fat condition, the contribution of lactate decreased to 13%, suggesting that fatty acid oxidation became a dominant source. This was supported by a study reporting that, in the perfused rat heart, rate values of triacyl glycerol degradation and total β-oxidation were markedly higher under high-fat and high-lactate conditions than low-fat and low-lactate conditions [24]. Plasma fatty acid levels have been shown to increase during exercise [25]. Thus, these findings may indicate that fatty acid oxidation in cardiac muscle is facilitated during intensive exercise, i.e., at high concentrations of blood lactate and free fatty acids. In the present study, LDH activity values in LV were markedly reduced in 30HT and 60HT groups (Fig 4B). This result may show that lactate oxidation in the heart was reduced and thereby the plasma lactate concentration may have been higher in these two groups than in the other groups during exercise. As mentioned above, in 60HT, but not 30HT, fatty acid metabolism was markedly improved in LV, thereby increasing the capacity of the heart to generate ATP via fatty acid oxidation during intensive exercise. Furthermore, CPT2 activity levels were remarkably up-regulated by 60HT in Gr and PL (Fig 3C). Thus, HT with daily hyperbaric exposure for 60 min has the potential to further increase exercise performance by facilitating fatty acid metabolism in skeletal as well as cardiac muscles.

Although significant changes were not observed, HT with hyperbaric exposure markedly upregulated DRP1 expression levels in LV (from 1.6-to 1.8-fold greater than HT group, Fig. 5A). DRP1 inhibition was shown to suppress mitochondrial respiration and reactive oxygen species without morphological changes in mitochondria, i.e. fission, in rat cardiomyocytes [26]. After acute exercise, mitochondrial fission was markedly observed in cardiac muscle [27]. In vascular smooth muscle cells, mitochondrial fission induced by platelet-derived growth factor was shown to enhance fatty acid oxidation and suppress glucose oxidation [28]. In the present study, expression levels of DRP1 were significantly correlated with both CS and HAD activity values in LV (Table 2). Thus, a modest increase in DRP1 levels may induce mitochondrial fission and contribute to promoting fatty acid metabolism in cardiac muscle.

In highly-trained mice as used in the present study, endurance training with hyperbaric exposure for 1 h markedly increased TFAM and DRP1 expression in Gr [2]. However, HT with hyperbaric exposure did not upregulate these protein expression levels (Fig 5). Thus, endurance exercise rather than HT may stimulate the expression of proteins concerning mitochondrial biogenesis in hind-leg muscles of highly-trained mice.

While hyperbaric exposure did not cause an additive effect, AKT1 expression levels (Fig 6B) were significantly increased in the four training groups in PL. A positive correlation was observed between AKT1 expression levels and FCSA values in PL (Table 2). Thus, hybrid training used in the present study caused muscle fiber hypertrophy, possibly via AKT1 upregulation. AKT1 was shown to promote muscle growth via facilitating satellite cell proliferation during skeletal muscle hypertrophy [8]. In PL, HSP70 expression levels were also significantly correlated with FCSA values (Table 2). In Gw, a significant correlation was observed between total FCSA and HSP70 expression levels, but not with AKT1 levels (Table 2). Mean values of HSP70 expression levels in Gw were markedly greater in 30HT (1.4-fold) and 60HT (1.6-fold) groups than in the HT group. This is consistent with previous findings using untrained [1] and highly-trained mice [2]. HSP70 was shown to play an important role in the recovery of striated muscle after severe exercise [29]. Recovery from muscle damage induced by daily exercise may be facilitated by hyperbaric exposure-induced HSP70 up-regulation, resulting in the promotion of muscular adaptation to exercise training.

Muscle capillary geometry was markedly improved by HT irrespective of hyperbaric exposure in PL (Fig 9). A positive correlation was noted between C:F ratios and AKT1 expression levels in PL (Table 2). In muscle-specific inducible AKT1 transgenic mice, the C:F ratio was significantly increased in the soleus muscle [30]. Thus, promoted expression of AKT1 by HT may contribute to improve muscle capillary network in highly-trained mice.

## Conclusions

The present study showed that hybrid training with daily hyperbaric exposure for longer than 30 min augmented exercise capacity in well-trained mice. Further studies on trained human subjects are needed to clarify whether exercise performance in athletes is promoted by hyperbaric exposure. In humans, middle ear barotrauma (MEB) was shown to be the most common side effect of hyperbaric oxygen therapy (approximately 3 ATA with 100% O_2_) [31, 32]. To reduce the risk of MEB, a compression rate may play an important role. The risk of MEB was shown to increase not only at a high rate of compression (0.28 ATA min^-1^) [31], but also at a low rate (0.068 ATA min^-1^) [32]. In the present study, the compression rate was set at 0.136 ATA min^-1^, which was shown to be the best rate for minimizing MEB and pain in humans [32]. However, commercial hyperbaric apparatuses currently compress air at a rate of 0.03-0.05 ATA min^-1^. To apply hyperbaric exposure to the daily training regimens of athletes, a new apparatus with a safer compression rate needs to be developed in the future.

## Acknowledgments

None

